# Unraveling the Link between Nutrition and Metabolic Syndrome Risk through In Silico Dietary Interventions

**DOI:** 10.1101/2024.07.04.602078

**Authors:** Drew Alessi, Chloe McCreery, Ali R. Zomorrodi

## Abstract

**Background:** Metabolic Syndrome (MetS) is a cluster of metabolic disorders that substantially increases the risk of chronic metabolic diseases. Diet is known to play a crucial role in the progression of MetS, yet a mechanistic understanding of its impact on MetS risk remains elusive.

**Methods:** To address this gap, we conducted a rigorous in silico diet intervention study by leveraging organ-resolved sex-specific whole-body models of metabolism. These models were utilized to computationally evaluate the effect of 12 diverse dietary regimens on key MetS biomarkers—glucose, triacylglycerol (TAG), LDL-C, and HDL-C—and fatty acid beta-oxidation in both males and females.

**Results:** Our analyses elucidated molecular mechanisms underlying the link between conventionally unhealthy diets and elevated MetS risk. Specifically, a typical Unhealthy diet indicated elevated TAG storage in the adipocytes and increased LDL-C to HDL-C ratios across both genders. Conversely, healthier dietary patterns like the Mediterranean and Vegan diets promoted favorable profiles for these biomarkers. Beyond substantiating these known dietary impacts, our analysis also revealed non-intuitive responses to diet. Notably, plant-based (Vegan and Vegetarian) diets induced elevated fatty acid oxidation compared to high-fat regimens like the Ketogenic diet, suggesting their potential in mitigating MetS risk. Pronounced gender differences in metabolic responses to diets were also observed, highlighting the need for gender-tailored dietary recommendations. Organ-specific dietary responses and their contributions to MetS biomarkers were also delineated, pinpointing the liver and lungs as major regulators of blood glucose homeostasis.

**Conclusions:** This study contributes to a deeper understanding of the intricate interactions between diet and MetS risk.

## Introduction

Metabolic Syndrome (MetS) is defined as a cluster of metabolic disorders that significantly increases the likelihood of developing several chronic diseases such as cardiovascular disease and other metabolic disorders, such as Type II Diabetes and Non-Alcoholic Fatty Liver Disease ^1^. MetS may involve any combination of metabolic disorders including hypertension (high blood pressure), hyperglycemia (high blood sugar), hyperlipidemia (high blood fat), hypercholesterolemia (high blood LDL-C), polycystic ovary syndrome (PCOS), insulin resistance, and excess body fat around the waist, otherwise known as central obesity ^1^. An individual suspected of MetS only needs to display symptoms from three of these metabolic diseases to be eligible for a diagnosis. It is important to note that MetS is not a disease in itself; rather, it is a cluster of interconnected metabolic abnormalities that constitute a pre-morbid condition, increasing the risk of developing other chronic metabolic diseases. MetS has been a growing problem in the Western world and across the globe ^2^. It is now estimated to affect approximately one-third of adults in the United States while its global prevalence ranges from 12.5% to 31.4%, depending on each region’s definition of MetS ^3^. The rise in MetS among the general population has significant implications for healthcare costs and the quality of life for affected individuals.

There is growing evidence that diet plays a significant role in developing MetS and is now considered a key modifiable risk factor. Previous studies suggest that diets high in added sugars, refined grains, and unhealthy fats can increase the risk of developing the condition ^4^. On the other hand, diets rich in whole grains, fruits, vegetables, and healthy fats protect against the development of MetS. For example, Mediterranean diet, well-balanced, or High Protein diet, are reported to be potentially advantageous for the management of MetS ^4^.

Due to the profound effect of diet on the risk of developing MetS, dietary interventions are considered as an attractive preventive and therapeutic approach for MetS. The common practice in existing dietary intervention studies is to assign one diet to each group of participants to isolate their effects on MetS. Nevertheless, the slowly developing nature of MetS makes it challenging to perform extensive dietary intervention studies.

Additionally, it is practically infeasible to examine the effect of multiple diets on human subjects due to difficulties associated with committing to various new dietary regimens particularly for long-term. Furthermore, since MetS affects many organs and metabolic processes, it is challenging to study and understand the effect of diet on the metabolic function of the many organs involved in maintaining metabolic homeostasis within the human body.

Computational studies are an intriguing alternative approach in digital medicine to investigate the effect of diet on human metabolism and metabolic disorders such as MetS. For example, a recent study developed a data-driven computational physiological model describing glucose, lipid, and cholesterol metabolism to investigate the metabolic changes that occur in response to a high-fat, high-sugar diet in mice ^5^. This study showed that this type of diet led to metabolic changes consistent with the development of MetS, including the increased production of fatty acids and inflammation. However, this model was limited only to a small proportion of the metabolism. Furthermore, such computational studies for humans, particularly at genome-scale, are still lacking.

Advances in the development of GEnome-scale Models (GEM) of metabolism have provided a promising route for computationally investigating the effect of diet on human metabolism at genome-scale. GEMs capture all metabolic reactions encoded by the genome of an organism. A series of global GEMs of human metabolism representing the metabolic capability of any human cell without specifying the organ-or cell-type-specific functions, have been reconstructed before ^6^. These models, which include Recon1 ^6^, Recon2 ^7^, Recon3D ^8^, and Human1 ^9^ have significantly enhanced our understanding of human metabolism. However, given that these models represent only the global human metabolism, they cannot capture organ-specific metabolic effects. Conversely, while several organ-, tissue-, and cell type-specific GEMs have been reconstructed ^10^, their scope is inherently limited to the metabolic pathways of their specified biological context. A paradigm-shift in this area was the development of two sex-specific organ-resolved whole-body models (WBMs) of human metabolism by Thiele et al ^11^. These models integrate GEMs for 20 organs, six sex organs, and six blood cell types into unified sex-specific models for male and female. The models have been parametrized using omics and physiological data and captures both whole-body metabolic processes and the effect of individual organs or cell types on the entire system. The applicability of this model has been demonstrated by integrating it with a dynamic coarse-gained mathematical model of glucose-insulin regulation to study disrupted metabolic processes in Type I Diabetes (T1D) at a whole-body scale ^12^. More recently, WBMs have been used to study host-microbiome interactions in Alzheimer’s disease ^13^ and host-virus co-metabolism during SARS-CoV-2 infection ^14^. They were also shown to accurately predict growth and biomarkers of inherited metabolic diseases in newborns and infants ^15^.

The WBMs offer an unprecedented opportunity to perform extensive in silico dietary intervention studies, taking into consideration organ-specific effects, yet this potential remains untapped in the realm of precision medicine. Here, we sought to use these WBMs to computationally evaluate the effect of a dozen diets on the risk of developing MetS and the role of different organs in this process. We simulated these dietary intakes in silico and evaluated their impact on organ-specific contributions to the serum levels of four key biomarkers of MetS, as well as the activity of the fatty acid oxidation pathway.

## Results

In this study, we used the organ-resolved whole-body GEMs of human metabolism for male and female ^11^ to computationally investigate the effect of diet on the risk of developing MetS. The male and female WBMs contain 81,094 and 83,521 metabolic reactions, respectively^11^. Here, we explored the effect of 12 different diets on the risk of developing MetS. These diets span a wide spectrum of dietary regimens and include an Average American diet, Average European diet, High Protein diet, High Fiber diet, Mediterranean diet, Vegetarian diet, Vegan diet, Gluten Free diet, DACH (Germany, Austria, and Switzerland) diet, Keto diet, a typical Balanced diet, and a typical Unhealthy diet. **Table 1** provides a brief description of each diet. The macronutrient and micronutrient breakdown of each diet are also shown in **Figure 1A** and **Supplementary File 1**, respectively.

**Figure 1.**
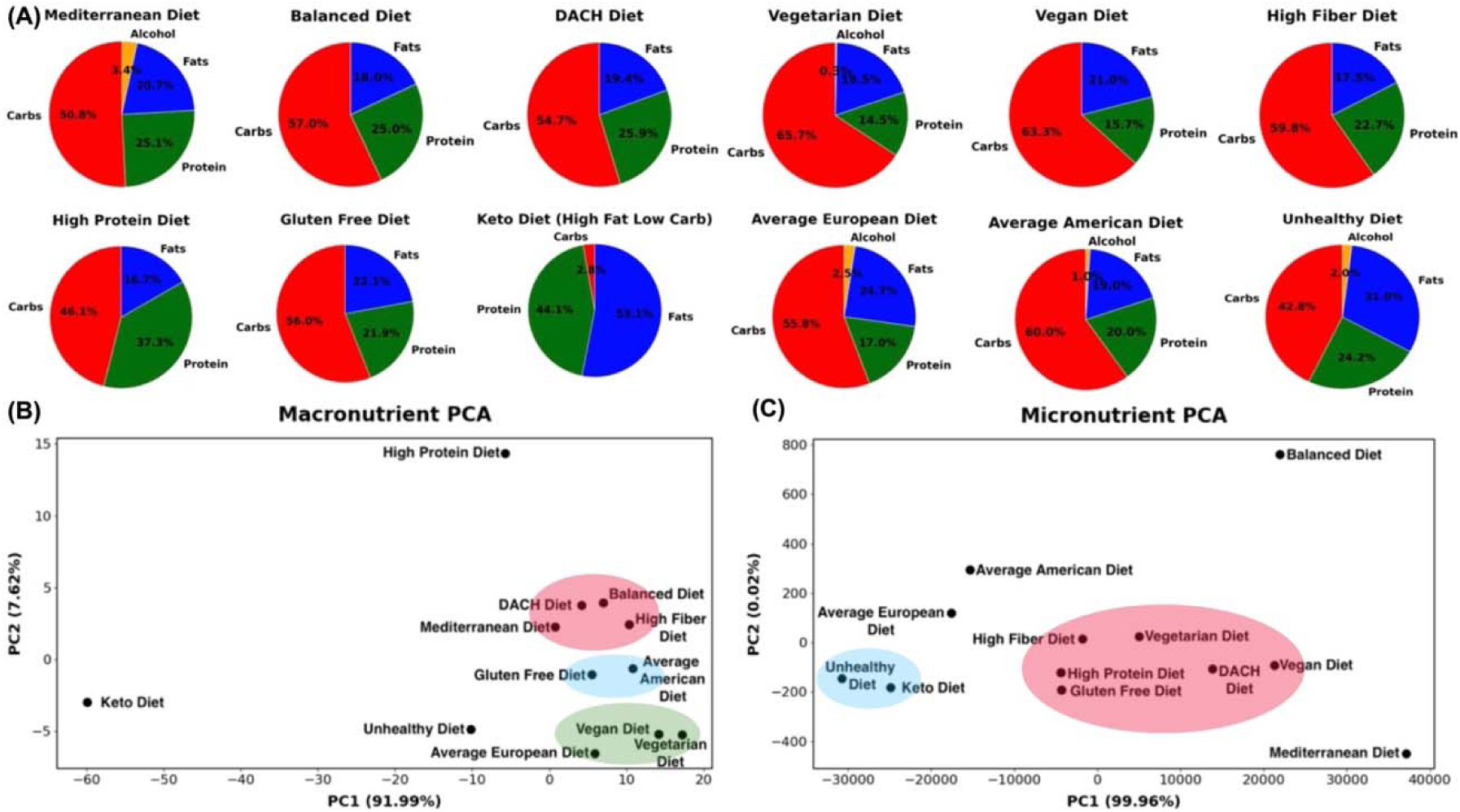
Macronutrient and micronutrient composition analysis of diets. (A) The macronutrient breakdown of the examined diets, (B) Spatial arrangement of diets based on their macronutrient, and (C) and micronutrient compositions. The diets were visualized using Principal Component Analysis (PCA), with reactions in the WBMs as features and diets as digital subjects (samples).

**Table 1.** Overview of the examined diets for this study.

| **Diet** | **Description** |
| --- | --- |
| Mediterranean Diet | The Mediterranean diet is made up of minimally processed foods, fresh fruits, and vegetables, and involves olive oil as its primary fat source. It assumes that dairy products are eaten daily, while poultry and fish are eaten occasionally, and red meat is eaten rarely. |
| Balanced Diet | The Balanced diet was designed to provide healthy amounts of essential nutrients, minerals, and vitamins to support normal metabolic functions for a healthy individual. |
| DACH Diet | The DACH diet, sourced from nutritional societies of Germany, Austria, and Switzerland. This diet was conceived to guarantee a nutritious health status for adults between the ages of 19 and 51 years old. |
| Vegetarian Diet | The Vegetarian diet is based on the most popular form of Vegetarianism, the lacto-ovo Vegetarian. This plant-based diet includes dairy and eggs but does not include any meat or fish. |
| Vegan Diet | The Vegan diet is another plant-based diet; however, it excludes all products derived from animals. |
| High Fiber Diet | The High Fiber diet contains animal sourced products and has a higher amount of dietary fiber than all other tested diets. |
| High Protein Diet | The High Protein diet is commonly used by athletes and those looking to lose weight. |
| Ketogenic (Keto) Diet | The Keto diet, also known as, the high fat low carb diet, aims to minimize the amount of carbohydrates intake and to replace them with fat and protein to promote the burning of body fat energy. |
| Gluten Free Diet | The Gluten Free diet excludes all foods containing gluten, a protein commonly found in wheat, rye, and barley. |
| Average European Diet | The components of this diet were based on an Austrian survey that was distributed in 2012 for approximately 1002 Austrian citizens across all age groups<sup>19</sup>. This diet entails moderate intake of fat, protein and alcohol, and low intake of carbohydrates and fiber. |
| Average American Diet | The American diet comprises high levels of processed foods, red and processed meat, refined grains, simple sugars, saturated fats, and sodium. |
| Unhealthy Diet | The typical Unhealthy diet consists of high intake of kilocalories, simple sugars, saturated fatty acids, and cholesterol as well as a low amount of dietary fibers often sourced from fruits and vegetables. |

To evaluate the effect of these diets on the risk of developing MetS, we mined prior knowledge on metabolic functions and molecular markers that are associated with the pathophysiology of MetS. This led us to identify four key molecular biomarkers directly implicated in MetS: glucose, triglycerides (also known as triacylglycerol or TAG), low-density lipoprotein cholesterol (LDL-C), and high-density lipoprotein cholesterol (HDL-C). We additionally chose to study the activity of the fatty acid beta-oxidation pathway. Although not a conventional marker, fatty acid oxidation plays a critical role in lipid metabolism, and the dysregulation of this pathway has been linked to MetS ^16–18^.

We evaluated how different diets may influence the serum levels of glucose, TAG, LDL-C, and HDL-C by analyzing reactions in the WBMs involved in their transport (exchange) between organs and the bloodstream. For the male WBM, we extracted 29 exchange reactions for glucose, three for TAG, three for HDL-C, and three for LDL-C. For the female WBM, we extracted 30 exchange reactions for glucose, three for TAG, three for HDL-C, and five for LDL-C. For the fatty acid beta-oxidation pathway, we focused on the last step of this process, which involves the breakdown of butyryl-CoA into acetyl-CoA in mitochondria and extracted two reactions from both the male and female WBMs representing this conversion in key tissues contributing to this pathway. The complete list of these reactions is provided in **Supplementary File 1.**

We performed separate in silico simulations for male and female using the sex-specific WBMs as typical male and female digital subjects. Computational simulations of the WBMs were performed using parsimonious FBA (pFBA) while constraining the whole-body maintenance reaction (see Methods).

### Analysis of diets’ nutrient composition and diversity

We analyzed the nutrient compositions of the 12 examined diets by employing Principal Component Analysis (PCA) using both their macro- and micronutrient compositions as features (**Figures 1B** and **1C**). Visualizing these diets using the first two principal components (PCs) revealed new insights into their similarities and dissimilarities based on underlying macro- and micronutrient compositions. The macronutrient PCA (**Figure 1B**) highlights a spectrum of dietary profiles with the Balanced, DACH, High Fiber, and Mediterranean diets clustering towards a balanced macronutrient composition, indicative of a harmonious blend of carbohydrates, proteins, and fats. Conversely, the Keto diet with an extreme fat preference and very low carbohydrates uptake, shows an extreme deviation for the rest of the diets. Another outlier is the High Protein diet, which heavily emphasizes animal-derived protein sources and diverges notably from other diets.

The Vegan and Vegetarian diets exhibited converging macronutrient compositions, emphasizing plant-based protein sources and a higher proportion of carbohydrates. The Average European diet is also positioned in proximity of the Vegetarian and Vegan diets, representing analogous proportions of carbohydrates and fats, coupled with a modest protein intake. The Average American and Gluten Free diets share similar profiles, suggesting comparable macronutrient distributions. The distinct positioning of the Average European diet in relation to its American counterpart reflects regional dietary preferences in Western diets. Finally, the Unhealthy diet’s distinct position indicates a skewed macronutrient profile.

The micronutrient PCA plot presents a different perspective on these diets, diverging from that for macronutrients (**Figure 1C**). The Balanced and Mediterranean diets are both distinctly separated from each other and from the remaining diets, indicating their unique micronutrient profiles. The High Fiber, Vegetarian, High Protein, DACH, Vegan, and Gluten Free diets aggregate into a pronounced cluster reflecting a congruence in their micronutrient compositions and potentially analogous impacts on metabolic processes and syndromes. The Unhealthy and Keto diets are clustered together on the left quadrant of the plot, representing their unique and similar micronutrient profiles that diverges from that of the healthier diets and may indicate potential nutrient deficiencies. In this landscape, the Average European diet is situated in the relative proximity of both the cluster of five diverse diets (including the High Fiber and Vegetarian), and the cluster of Unhealthy and Keto diets, suggesting a partial overlap in micronutrient profiles with both groups. In contrast, the Average American diet remains further from both clusters, implying a potentially distinct micronutrient profile that does not align with either of these diet clusters or with its European counterpart. Collectively, these analyses indicate the marked diversity in nutritional content across these diets. s

### The effect of diet on the systemic metabolic response

We examined how diverse dietary patterns influence the systemic metabolic response in males and females. To this end, we utilized t-distributed Stochastic Neighbor Embedding (t-SNE) analysis with all the 81,094 and 83,521 metabolic reactions within the male and female WBMs, respectively. In these analyses, reactions served as features while the 12 diets served as samples with each diet corresponding to a digital subject on the respective diet. This analysis revealed the differential impact of these diets on the systemic metabolic response (**Figure 2A** and **2B**). A discernible pattern is that the diets traditionally known to be healthier, including the Mediterranean, Balanced, Vegan, and DACH diets, manifest as a cluster in both males and females, reflecting their analogous and potentially beneficial modulation of metabolic state. Conversely, the Unhealthy and Keto diets are conspicuously isolated from these healthy diets and positioned on the opposing corners of the plots, implying a divergent influence on the systemic metabolism. The Vegetarian diet, while positioned separately, tends towards the healthier diets cluster in male and female, suggesting a favorable metabolic imprint akin to these diets. The High Protein, High Fiber, and Gluten Free diets occupy a middle locus in both males and females, which indicates a rather moderate influence on metabolic landscape, neither heavily favoring nor significantly diverging from a state of metabolic health. We also observe a notable dispersion of diets within these clusters, indicating a subtle gradation in their metabolic influence. Intriguingly, the Average American diet is clustered with the Average European diet and is located near the cluster of less healthy (Unhealthy and Keto) diets in males; however, it moves to a distinct position near the cluster of healthier (Vegetarian, DACH, Balanced) diets in females.

**Figure 2.**
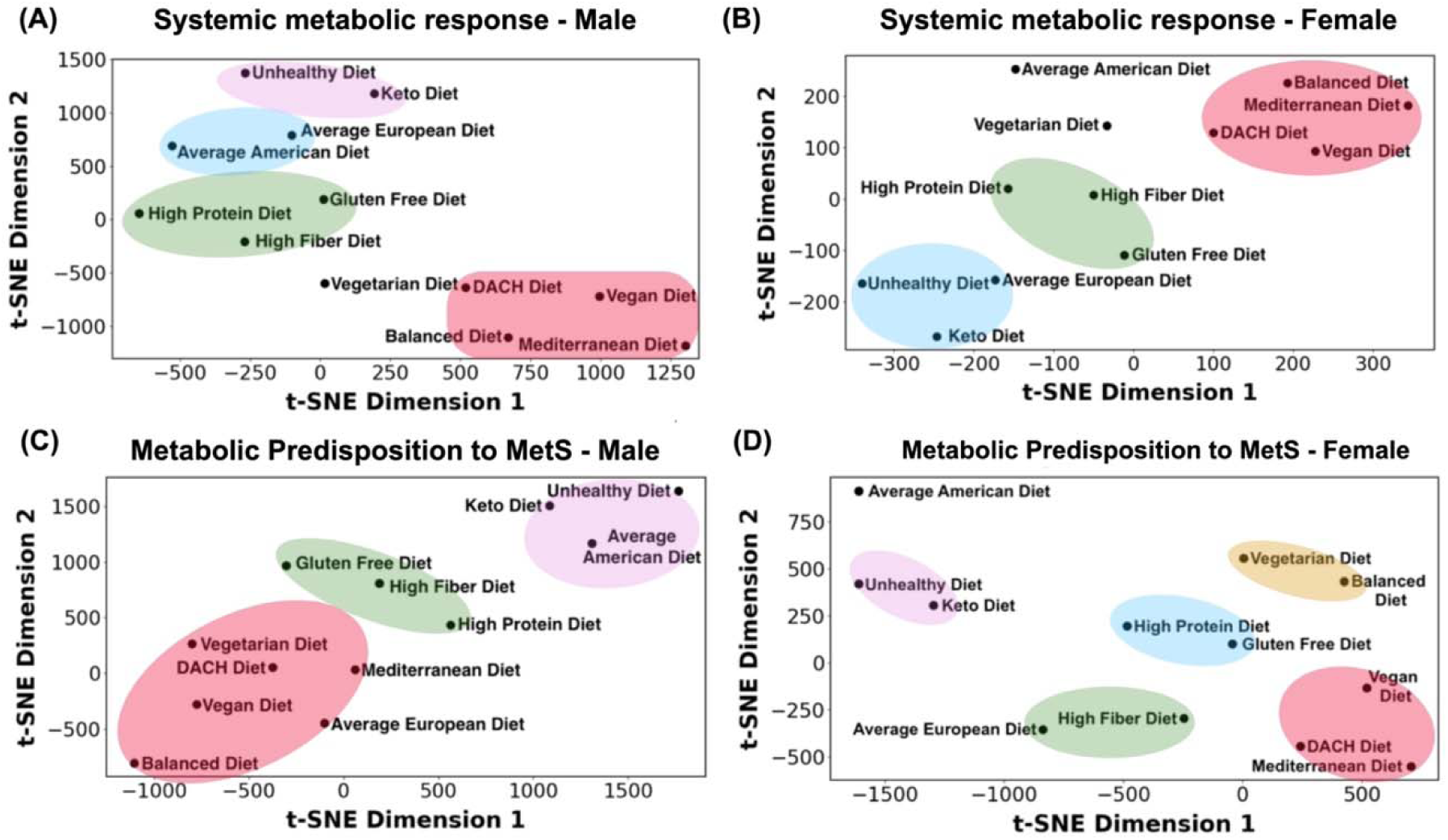
Visualization of the examined diets based on the systemic metabolic response and metabolic predisposition to MetS. (A) Systemic metabolic response for the male and (B) female WBMs. (C) Metabolic predisposition to MetS for the male and (D) female WBMs. The diets were visualized using t-Distributed Stochastic Neighbor Embedding (t-SNE), with reactions in the WBMs as features and diets as digital subjects (samples).

### The effect of diet on metabolic predisposition to MetS

To evaluate the interplay between diet and susceptibility toward developing MetS, we leveraged reactions in the male and female WBMs responsible for glucose, TAG, HDL-C, and LDL-C exchange with blood, as well as the reactions involved in fatty acid oxidation. These reactions were used as features for visualization using t-SNE with the 12 diets serving as digital subjects (**Figures 2C** and **2D**). The t-SNE plot in males depicted a continuum of dietary impacts on metabolic processes pertinent to MetS with trends somewhat like those seen in **Figure 2A**, where healthier and unhealthy diets are clustered distinctly separated in the opposing corners of the t-SNE plot and the remaining diets positioned between these two extremes. A noticeable divergence between the two is the repositioning of the Average European diet from the proximity of unhealthy diets in **Figure 2A** to the cluster of conventionally healthy diets in **Figure 2C**. This suggests a MetS risk profile for this diet similar to that of the healthy diets.

The t-SNE plot for females presents a rather distinct clustering of diets in relation to the MetS metabolic biomarkers compared to males (**Figure 2D**). A notable aggregation is evident in the lower right quadrant of the female t-SNE plot, comprising the Mediterranean, Vegan, and DACH diets. However, the other two healthy diets, namely the Balanced and Vegetarian diets, form an independent cluster in the upper right quadrant, representing a differentiated MetS risk profile compared to other healthy diets in females. While closer to the Unhealthy and Keto diets, the Average American diet is markedly set apart in the upper right corner, hinting at a unique MetS risk profile induced by this diet in females. Its European counterpart, however, along with the High-Fiber diet, is positioned closer to the cluster of healthier diets, similar to that in males. These observations underscore the specific metabolic response of females to both healthy and unhealthy diets in relation to the MetS biomarkers.

### The effect of diet on individual MetS biomarkers

In the subsequent sections, we evaluate the effect of diet on each molecular biomarker of MetS and fatty acid oxidation as well as the significance of different organs/tissues.

#### Glucose

Glucose is one of the most widely recognized diagnostic markers for MetS ^20^. Specifically, abnormally elevated fasting serum levels of glucose are considered a major MetS risk factor. We assessed a total of 24 reactions in males and 31 reactions in females involved in the transport of glucose between different organs/tissues and the systemic blood circulation.

##### Overall glucose secretion into the blood

Evaluating the total glucose secretion into the bloodstream (i.e., sum of glucose secretion fluxes by all organs secreting glucose) in the male WBM revealed that surprisingly the Balanced diet has the highest glucose secretion flux (3,994.7 mmol/person/day), while the Unhealthy diet (3,266.3 mmol/person/day) followed the Keto diet (3,728.9 mmol/person/day) and Average American diet (3,760.4 mmol/person/day) exhibit the lowest glucose section fluxes (**Figure 3A**). The rest of the diets displayed a consistent overall glucose secretion of 3,905.4 mmol/person/day in the male WBM. Similar patterns were observed in the female WBM, where, contrary to expectations, the healthier diets show the highest overall glucose secretion into the blood, while the unhealthy diet shows the reduced glucose secretion fluxes. Specifically, the Mediterranean diet shows the highest overall glucose secretion flux (7,228.0 mmol/person/day), followed by the Vegan (7,165.4 mmol/person/day), Balanced (7,161.0 mmol/person/day), and DACH (7,157.6 mmol/person/day) diets (**Figure 3B**).

**Figure 3.**
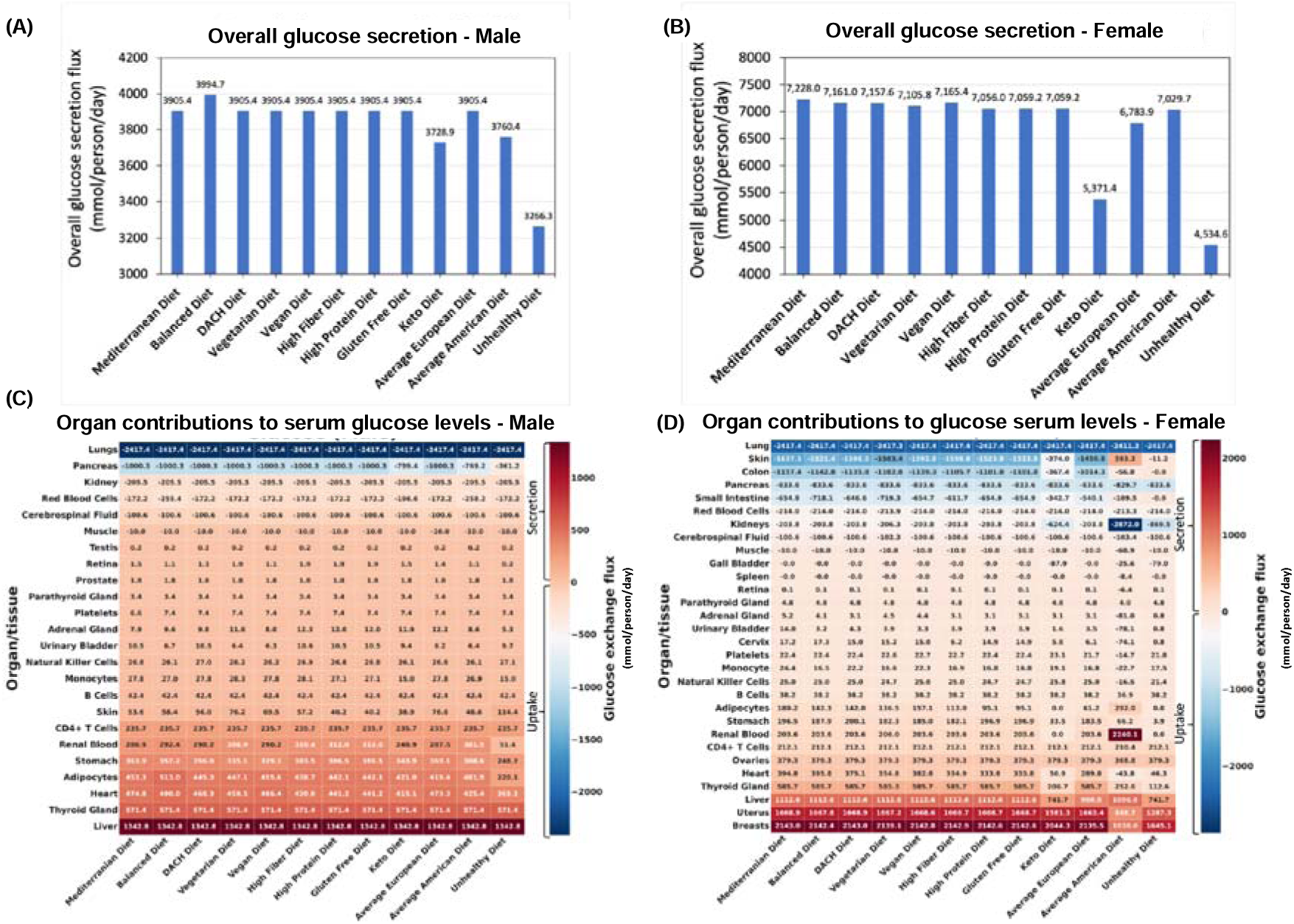
Impact of diet on blood glucose levels. (A) Predicted overall glucose secretion flux into the bloodstream for the male and (B) female WBMs. (C) Contribution of different organs/tissues within the male and (D) female WBMs to glucose secretion and uptake into/from the blood.

Conversely, the Unhealthy diet exhibits a significantly lower overall glucose secretion flux compared to all other diets (4,534.6 mmol/person/day). The Keto diet (5,371.4 mmol/person/day) and Average European (6,783.9 mmol/person/day) diets also revealed particularly low secretion levels similar to those in males.

##### Organs/tissues secreting glucose into the blood

Our analysis identified 18 organs/tissues in both male and female WBMs that consistently engaged in glucose secretion into the bloodstream (**Figure 3C** and **3D**). The liver, thyroid gland, heart, adipocytes, and stomach emerged as major contributors to glucose secretion into blood in the male WBM. In the female WBM, the breast, uterus, and liver emerged as the organs with the highest contribution to glucose secretion into blood across all diets (**Figure 3D**).

##### Organs/tissues taking up glucose from the blood

Six organs/tissues in male and 11 in female WBMs demonstrated consistent glucose uptake from blood across all the diets. In males, the lungs emerged as the most avid consumer with a consistent glucose uptake flux of 2,417.4 mmol/day/person across all studied diets, followed by pancreas (1,000.3 mmol/day/person), and kidney (205.5 mmol/day/person) **(Figure 3C).** In females, the lung, skin, and colon are responsible for taking up the highest levels of glucose from the blood across most diets (**Figure 3D**).

##### Organs with notable altered contributions to glucose serum levels across diets

The pancreas, adipocytes, and renal blood are among the organs/tissues in males for which their contributions to serum glucose levels are most influenced by diet variations **(Figure 3C).** The diets that induced the largest change in glucose secretion or uptake by these organs is the Unhealthy diet, followed by the Average American and Keto diets.

Notably, these are the three diets that clustered together with respect to MetS risk (**Figure 2C**). For instance, the glucose uptake flux from the blood by the pancreas recorded to be 361.2, 769.2, and 799.4 mmol/day/person for the Unhealthy, Average American, and Keto diets, respectively, which are significantly lower than that for all other diets (1,000.3 mmol/day/person). Adipocytes, which act as the WBM’s representation of fat storage distributed throughout the body, also experienced marked changes in glucose secretion in the male WBM under the Unhealthy, Average American, and Balanced, diets. The Balanced (513.0 mmol/day/person) diets show a notable increase in adipocytes glucose secretion flux into the blood relative to other diets (**Figure 3C**). Conversely, the Unhealthy diet followed by the Average American diet resulted in a substantial decrease in adipocytes’ glucose secretion (220.1 and 401.9 mmol/day/person, respectively). The renal blood was another organ with altered glucose secretion profile across diets in the male WBM. In the WBMs, the kidney is associated with two distinct reactions involving the exchange of glucose with blood: the exchange of glucose between the kidney and systemic blood, and the exchange of glucose between the renal blood and systemic blood (depicted by ‘Kidney’ and ‘Renal Blood’, respectively in **Figure 3C**). Notably, the kidney is involved in glucose uptake from the systemic blood circulation, while the renal blood engages in glucose secretion into the systemic blood circulation in both the male and female WBMs. This can be explained by previous reports that the kidneys play a significant role in maintaining glucose homeostasis in the body by releasing glucose into the circulation via gluconeogenesis, taking up glucose from the circulation to fulfill their energy needs, and reabsorption of glucose at the proximal tubule ^21^. Here, we observed an increase in glucose secretion flux from renal blood into systemic blood in the male WBM under the High Protein and Gluten Free diets (312.0 mmol/day/person for both) as well as the High Fiber diet (310.4 mmol/day/person). In contrast, a significant decrease was observed under the Unhealthy diet (51.37 mmol/day/person) compared to the remaining nine diets (300.9 ± 18.5 mmol/day/person) **(Figure 3C)**. No changes were observed under these diets for glucose uptake from the systemic blood circulation by the kidney.

In females, the organs that are most affected by diet include the breast, uterus, liver, skin, and colon (**Figure 3D**). Specifically, the glucose secretion fluxes for the breast and uterus under the Average American and Unhealthy diets were significantly lower compared to other diets. Similarly, the liver exhibits a notable decrease in glucose secretion flux under the Unhealthy, Keto, and Average European diets compared to other diets—a pattern not observed in the male WBM. Skin and colon also experience diminished glucose uptake from the blood under the Unhealthy, Average American, and Keto diets. Notably, we noticed several outlier responses for glucose under the Average American diet for the female WBM. For example, while skin cells engage in glucose uptake from blood under all other diets, they show glucose secretion rather than uptake under the Average American diet. Similar abnormalities were observed in the retina, adrenal gland, urinary bladder, cervix, platelets, monocytes, NK cells, and the heart.

#### TAG

Formed by the esterification of three fatty acid molecules to glycerol, TAG is a prevalent form of fat and primary energy storage within the human body. Elevated serum levels of TAG are a hallmark of MetS ^20, 22^. Elevated TAG serum levels are often associated with the consumption of a high-calorie diet rich in refined carbohydrates and saturated fats, which can contribute to dyslipidemia– another leading factor implicated in MetS ^4^. TAG biosynthesis occurs predominantly in the liver, where excess dietary carbohydrates and proteins—surpassing the body’s immediate needs—are converted into fatty acids and subsequently into TAG. Concurrently, the small intestine is instrumental in the absorption of dietary fats, predominantly in the form of TAG. Before entering the bloodstream, TAG is packaged into very low-density lipoproteins (VLDL) in the liver and chylomicrons in the intestine as it is not soluble in the aqueous environment of the blood. Adipose tissues serve as the principal depots for TAG storage. Upon arrival via the bloodstream, TAG is hydrolyzed back into free fatty acids and glycerol, mediated by the enzyme lipase. The liberated fatty acids and glycerol are then absorbed by adipocytes and re-esterified into TAG for long-term storage ^22^.

In our analyses, we assessed TAG storage within adipocytes in the WBMs as a proxy for the TAG serum level. This was accomplished by examining the flux of three specific TAG synthesis reactions in adipocytes within both the male and female WBMs, each responsible for the esterification of different types of essential and non-essential fatty acids into TAG within the adipocytes. We computed the aggregate flux through these three reactions, treating this sum as a representation of total TAG synthesis and accumulation within adipocytes.

Our analysis revealed a pronounced disparity in TAG synthesis and storage within adipocytes across various diets in both male and female WBMs. In males, the Unhealthy diet exhibits a considerably higher TAG synthesis flux of 21.61 mmol/day/person compared to remaining diets (1.86 ± 0.92 mmol/day/person) **(Figure 4A)**, which can be attributed to the diet’s excessive calorie load and unhealthy fats intake. Conversely, the Balanced diet is characterized by a negligible TAG synthesis, suggesting a dietary composition that mitigates excessive lipid accumulation. Contrary to expectation, the Average American Diet, despite its reputation for high caloric content and associations with MetS, also shows negligible TAG synthesis in adipocytes. The rest of the diets show comparable TAG synthesis in adipocytes although the Vegetarian diet exhibits slightly lower TAG synthesis flux (1.99 mmol/day/person) relative to remaining diets.

**Figure 4.**
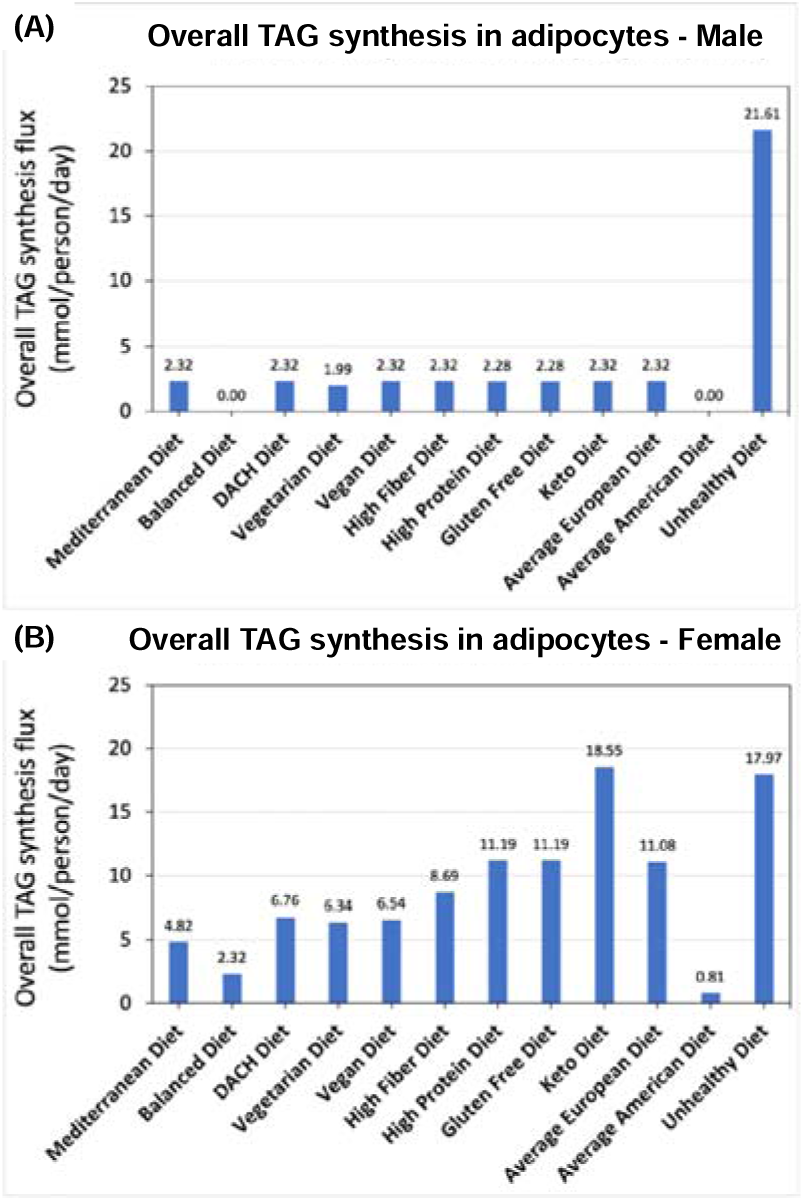
Impact of diet on TAG synthesis. (A) Predicted overall TAG synthesis in the adipocytes in the male and (B) female WBMs.

Our analysis using the female WBM revealed that TAG synthesis and storage in adipocytes was highest under the Keto (18.55 mmol/person/day) and Unhealthy diets (17.97 mmol/person/day) **(Figure 4B)**. The high fat content of the Keto diet (more than 53%; **Figure 1**) appears to significantly increase the TAG synthesis in females, a pattern diverging from that for the males. Again, consistent with males but counterintuitively, the Average American shows the lowest TAG synthesis in females (0.81 mmol/person/day), The next diets with low TAG synthesis and storage in adipocytes are the Balanced (2.32 mmol/person/day) and Mediterranean (4.82 mmol/person/day) diets, aligning with our expectation for these healthier diets.

#### LDL-C and HDL-C

HDL-C and LDL-C are two types of lipoproteins that transport cholesterol throughout the body via the blood circulation thereby regulating cholesterol levels. LDL-C (often referred to as “bad cholesterol”) is responsible for transporting cholesterol from the liver to various tissues within the body ^23^. HDL-C (also known as the “good cholesterol”) engages in reverse cholesterol transport, removing excess cholesterol from the bloodstream and peripheral tissues and transporting it back to the liver for disposal. Elevated levels of LDL-C can lead to the build-up of cholesterol in the arteries, forming plaques that can narrow down and obstruct blood vessels—a condition referred to as atherosclerosis, which increases the risk of MetS and cardiovascular disease.

Conversely, high HDL-C serum levels, have a protective effect against atherosclerosis ^24^, while its low serum levels contribute to the increased prevalence of MetS, particularly in males ^25^.

In our analyses, we focused on three distinct reactions for HDL-C and three others for LDL-C within the male WBM that contribute non-zero (although sometimes negligible) flux to HDL-C and LDL-C exchange with blood within the WBMs—namely the liver, muscle, and adipocytes. For the female WBM, we selected three reactions for HDL-C and five reactions for LDL-C that exchange HDL-C and LDL-C between the bloodstream and liver, muscle, adipocytes, kidney, and renal blood.

##### Overall LDL-C/HDL-C blood secretion ratio

We first evaluated the total secretion flux of HDL-C and LDL-C by all relevant organs for each diet and then calculated the ratio of total LDL-C to HDL-C secretion fluxes (LDL-C/HDL-C). For the male WBM, this analysis revealed that the Unhealthy diet exhibits a substantially higher LDL-C/HDL-C ratio (1.54) compared to all other diets (**Figure 5A**), indicating impaired cholesterol metabolism and potentially increased risk for MetS. The ratios for the rest of the diets are comparable and very close to 1, although there are slight variations (0.98 ± 0.01). The Average American diet shows the second highest LDL-C/HDL-C ratio (1.01), which underscores the diet’s association with adverse lipid profiles and heightened MetS risk. In contrast, the Vegan diet shows the lowest LDL-C/HDL-C ratio (0.97) in males, indicating a more favorable lipoprotein profile. The remaining diets exhibit intermediate LDL-C/HDL-C ratios, reflecting a moderate influence of these diets on cholesterol metabolism.

**Figure 5.**
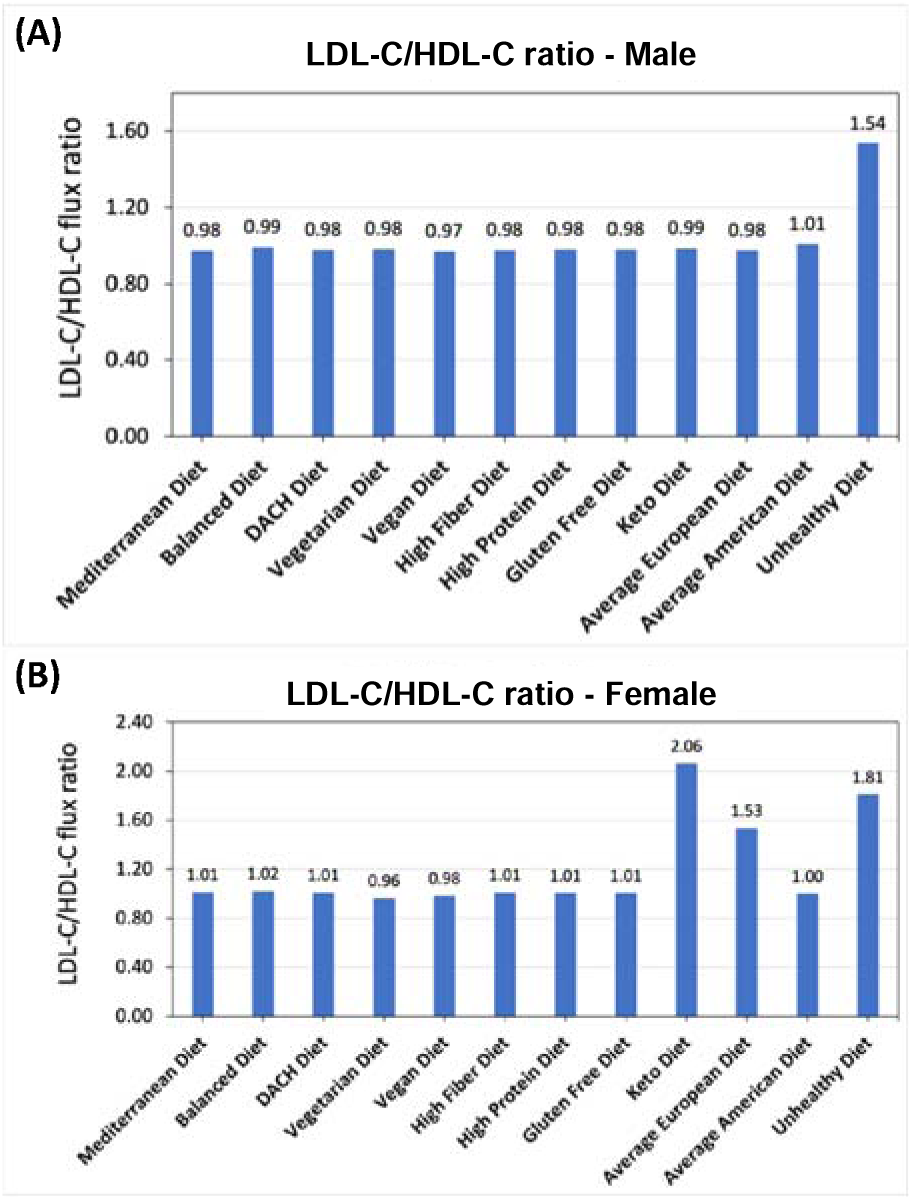
Effect of diet on LDL-C to HDL-C ratio. (A) Predicted LDL-C/HDL-C ratio in the male and (B) female WBMs.

When examining the LDL-C/HDL-C ratio in females, more variations were observed in response to diet compared to males. The greatest ratios were noted under the Keto (2.06) and Unhealthy (1.81), and Average European (1.53) diets. Conversely, the lowest LDL-C/HDL-C ratios occurred under the Vegetarian (0.96) and Vegan (0.98) diets **(Figure 6B)**. The remaining diets (Mediterranean, Balanced, DACH, High Fiber, High Protein, Gluten Free, and Average American diets) revealed intermediate ratios ranging from 1.00 to 1.02. Again, we expected a higher LDL-C/HDL-C ratio for the Average American diet, considering its high intake of saturated fats.

**Figure 6.**
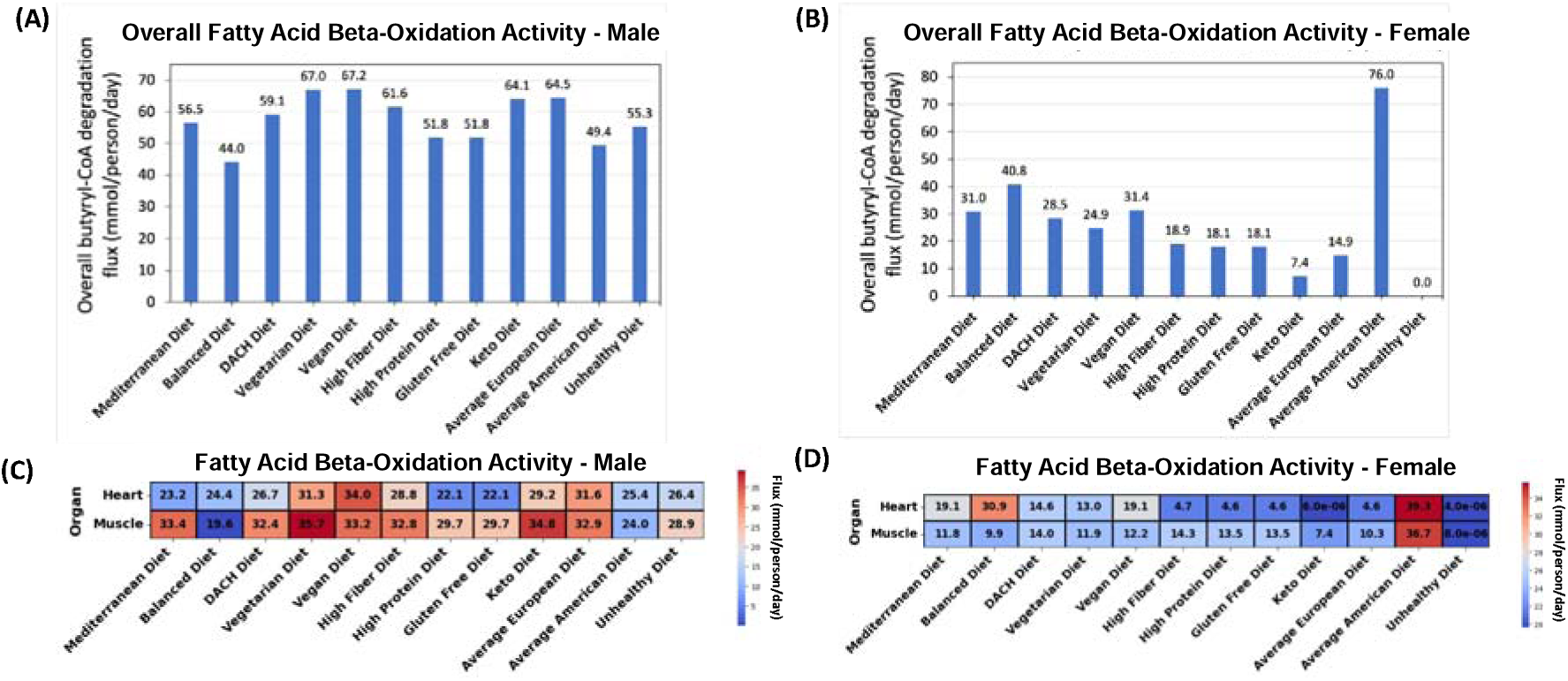
Effect of diet on the fatty acid beta-oxidation pathway. (A) Predicted overall fatty acid oxidation activity for the male and (B) female WBMs. (C) Contribution of the (skeletal) muscles and heart to fatty acid oxidation in the male and (D) female WBMs.

#### Fatty acid beta-oxidation pathway

Fatty acid beta-oxidation pathway plays a pivotal role in the human body’s energy metabolism by breaking down fatty acids to generate ATP. This pathway’s efficiency is crucial for maintaining metabolic health as deficiencies can lead to elevated free fatty acid levels, which contributes to metabolic diseases including MetS ^26, 27^. However, the complexity of this metabolic process and its limited experimental accessibility in human cells has hampered a deep understanding of the pathophysiological mechanisms underlying the link between dietary factors, dysregulation of this pathway, and the risk of metabolic diseases, especially MetS ^26, 27^.

We explored the effect of various diets on the activity of fatty acid beta-oxidation pathway. Our analysis centered on the final step of this metabolic process, the conversion of butyryl-CoA, a 4-carbon fatty acid, into acetyl-CoA, a 2-carbon molecule, catalyzed by the enzyme thiolase. The flux of this reaction was used as a surrogate for the activity of the fatty acid oxidation process. Although fatty acid beta-oxidation occurs in all tissues and organs within the human body, the (skeletal) muscles and heart (cardiac muscles) are the major contributors ^28^. Therefore, we focused our investigations especially on the muscles and heart in both the male and female WBMs. This rationale guided us to select two specific reactions within the male and female WBM representing butyryl-CoA breakdown in these tissues/organs.

##### Overall fatty acid beta-oxidation activity

We first evaluated the overall fatty acid beta-oxidation activity within the human body as captured by sum of the butyryl-CoA breakdown fluxes in the muscles and heart. For the male WBM, the Vegan (67.2 mmol/person/day) and Vegetarian (67.0 mmol/person/day) diets followed by the Average European (64.5 mmol/person/day) and Keto (64.1 mmol/person/day) diets exhibited the highest overall activity (**Figure 6A**). On the other end of the spectrum, unexpectedly the Balanced diet followed by the Average American diets showed the lowest activity (44.0 and 49.4 mmol/person/day, respectively). The remaining diets show intermediate activity.

For the female WBM, a substantially higher activity of fatty acid beta-oxidation was observed for the Average American diet (76.0 mmol/person/day) compared to other diets (**Figure 6B**)–a pattern contrary to the male WBM. The Balanced (40.8 mmol/person/day Vegan (31.4 mmol/person/day), Mediterranean (31.0 mmol/person/day), and DACH (28.5 mmol/person/day) diets are the next diets high fatty acid oxidation activity. Conversely, the Unhealthy and, unexpectedly, the Keto diets demonstrated the lowest activity (0.0 and 7.4 mmol/person/day, respectively).

##### The effect of diets on organs’ contributions to fatty acid oxidation

In the male WBM, the highest fatty acid oxidation activity in muscles was observed under the Vegetarian diet and Keto diets (35.7 and 34.8 mmol/person/day, respectively; **Figure 6C)**. The lowest activity in muscle was observed for the Balanced (19.6 mmol/person/day) followed by the Average American (24.0 mmol/person/day) and Unhealthy (28.9. mmol/person/day) diets. Similar observations were made for the heart which, like muscle, depends on fatty acids as a primary energy source—a process is crucial for the heart’s constant contraction and efficient functioning ^29^. For the heat, the Vegan, Average European, and Vegetarian diets exhibited the highest activity (34.0, 31.6, and 31.3 mmol/person/day, respectively). Conversely, the High Protein and Gluten Free diets (22.1 mmol/person/day for both) exhibited the lowest activities.

For the female WBM, the Average American diet induces a substantially higher fatty acid oxidation activity in both the muscles (36.7 mmol/person/day) and heart (39.3 mmol/person/day) compared to other diets (**Figures 6D**). The High Fiber (14.3 mmol/person/day) and DACH (14.0 mmol/person/day) diets exhibit the next highest activity in the muscles. This is supported by existing literature that dietary fiber can induce metabolic adaptations in the muscles, potentially upregulating the activity of fatty acid oxidation ^30,31^. For the heart, the Balanced, Mediterranean, and Vegan diets lead to the next highest fatty acid oxidation activity (30.9, 19.1, and 19.1 mmol/person/day, respectively). Conversely, the lowest activity in both the muscles and heart was observed for the Unhealthy and Keto diets.

## Discussion

We systematically evaluated the impact of 12 diverse dietary regimens on key biomarkers associated with MetS through the lens of sophisticated, sex-specific, organ-resolved whole-body models of metabolism. Our investigation explored the secretion of glucose, TAG, LDL-C, and HDL-C into the bloodstream, as well as fatty acid oxidation activity in both male and female. This in silico analysis revealed how specific dietary patterns influence the physiological processes underlying these MetS biomarkers and uncovered new insights into the contribution of different organs and tissues.

Our study supports the existing hypotheses that unhealthy diets—characterized by low intake of dietary fibers and high intake of saturated fats, cholesterol, and simple sugars— tend to adversely impact MetS biomarkers, while healthy diets, i.e., those involving the intake of whole grains, fruits, vegetables, lean proteins, and healthy fats promote metabolic homeostasis and reduce MetS risk. Nonetheless, we also identified counterintuitive relationships between specific diets and MetS biomarkers, indicating the complex interplay between dietary intake and physiological processes underlying MetS based on the WBMs.

### The analysis of diets based on nutrient composition revealed significant variability and highlights the importance of considering both macro and micronutrient profiles

PCA provided a macroscopic view of the dietary patterns examined in our study, revealing distinct clusters of diets based on their macronutrient **(Figure 1B)** and micronutrient compositions **(Figure 1C)**. This analysis illustrated a gradient of macro and micronutrient diversity and density in the examined diets, with distinct spatial distribution of diets in the micronutrient PCA plot compared to macronutrients. Specifically, despite both being considered healthy diets, the Mediterranean and Balanced diets occupy conspicuously isolated positions in the micronutrient PCA plot, separate from each other and from the remaining diets—a pattern not observed in the macronutrient PCA plot.

This suggests a unique micronutrient profile for these diets, diverging from the remaining diets. The Mediterranean diet, rich in plant-based foods, olive oil, and fish, likely provides a distinct array of vitamins, minerals, and phytonutrients. The Balanced diet’s unique positioning may also reflect its diverse and equilibrated nutrient composition that adheres to nutritional guidelines for promoting metabolic health. The Average American diet also stands apart from the cluster of unhealthy diets in the micronutrient PCA plot, emphasizing its unique micronutrient composition. These observations highlight the importance of considering both macro and micronutrient compositions when investigating the effect of dietary regimens on metabolic health.

### Examining the impact of diet on systemic metabolic responses and MetS predisposition revealed clear separations between healthy and unhealthy dietary patterns across both genders

Our t-SNE analysis offered a deeper dive into the effects of these diets on both systemic metabolic response **(Figures 2A** and **2B)** and metabolic predisposition to MetS (**Figures 2C** and **2D**) in male and female. Analysis of metabolic response at the systemic level revealed that diets traditionally considered healthy—such as the Mediterranean, Vegan, DACH, and Balanced—formed clusters that significantly diverged from those deemed less healthy, like the Unhealthy and Keto diets, in both male and female. A similar divergence of the healthy and unhealthy diet clusters was also observed in the t-SNE plots representing metabolic susceptibility to MetS. This spatial stratification highlights the notably distinct impacts of these dietary groups on both the systematic metabolic homeostasis and MetS. It also demonstrates the marked potential of diet to modulate the biochemical milieu associated MetS. The consistent clustering of the Keto diet with the Unhealthy diet suggests potential adverse metabolic outcomes despite its popularity for weight loss. In addition to these commonalities between the systematic metabolic response and MetS susceptibility, we noticed differences between the two as well. A notable deviation between the two is the repositioning of the Average European diet from the close proximity of the Unhealthy diet when examining systematic metabolic response **(Figures 2A** and **2B**) to that of the cluster of healthier diets when exploring response to MetS risk (**Figures 2C** and **2D**) in both the male and female WBMs. This indicates a MetS risk profile of this diet that is closer to the healthier diets, which is in contrast to its American counterpart showing close proximity to unhealthy diets with respect to both systematic metabolic response and MetS risk.

### Unhealthy diets yield paradoxically lower glucose secretion profiles despite higher glycemic loads compared to healthier diets

Analyzing the impacts of diet on individual MetS biomarkers using the WBMs provided granular insights into the complex interactions between dietary intake and the physiological indicators. For instance, when examining the effect of diet on glucose secretion into the systemic blood, the Unhealthy, Keto, and Average American, diets resulted in lower overall glucose secretion levels into the blood compared to healthier diets in both the male and female WBMs (**Figures 3A** and **3B**). Specifically, the Balanced and the Unhealthy diets showed the highest and lowest glucose secretion, respectively, in males. Similar patterns were observed in females, where the Mediterranean diet led to the highest glucose secretion and the Unhealthy diet showed the lowest. These results contrast with our expectations that unhealthy diets are presumed to elevate glucose levels due to higher glycemic loads. This counterintuitive finding might be attributed to the body’s complex metabolic processes and its ability to maintain homeostasis. For instance, the body might be compensating for the high intake of simple sugars in unhealthy diets by reducing glucose secretion. Additionally, the composition of these unhealthy diets, often high in simple sugars, could cause a rapid spike in blood glucose levels followed by a sharp decline, leading to lower overall glucose secretion.

Another seemingly paradoxical observation is that despite significantly lower carbohydrate content in the Keto diet, it shows higher glucose secretion levels into the blood relative to the Unhealthy diet in both the male and female WBMs. This observation challenges the simplistic notion that lower dietary carbohydrate intake directly translates to reduced blood glucose. The Keto diet, with its high fat content, is known to fuel gluconeogenesis, the production of glucose from non-carbohydrate sources ^32, 33^, which may contribute to the observed glucose secretion profiles compared to the Unhealthy diet. These findings could be also influenced by the inherent limitations of the WBMs.

Importantly, these observations suggest that the risk of MetS may not solely be determined by the healthiness of a diet or its carbohydrate contents but also by how the body metabolically responds to different dietary patterns. Further studies are needed to validate these computational findings.

### Organ-specific contributions to blood glucose homeostasis reveal the critical role of the liver and lungs, highlighting their pivotal influence on MetS risk

The organs identified to play a major role in blood glucose levels align with established physiological knowledge. For instance, the liver, a known site for gluconeogenesis and glycogenolysis, emerged as a major glucose secreting organ in both males and females (**Figures 3C** and **3D**). Our study also identified the lungs as a major contributor to glucose blood levels, exhibiting the highest glucose uptake from the blood in both males and females, although the uptake levels remained unaffected by dietary variations. This major role of the lungs in glucose metabolism is consistent with previous reports ^34^ and implies that any impairments in the lung function can critically affect serum glucose levels and, therefore, raise the risk of developing MetS. This finding is supported by existing evidence that lung conditions such as asthma and pulmonary hypertension are associated with abnormally high postprandial blood glucose levels ^35^. In addition, Baffi et al. ^36^ have discussed potential mechanisms that explain the associations between MetS and lung health. These highlight that the lung’s role in managing the blood glucose homeostasis and MetS risk is crucial yet often overlooked.

### Dietary variations, especially diets high in fats and simple sugars, markedly influence blood glucose contributions from renal blood, pancreas, and adipocytes in males, and from breasts, uterus, and liver in females

Interestingly, our findings revealed that specific organs are more responsive to dietary changes than others. In males, the renal blood, pancreas, and adipocytes exhibited the most significant response to dietary variations. In particular, an increase in glucose secretion from renal blood into the systemic blood was observed under the High Protein, Gluten Free, and High-Fiber diets and a notable decrease was noted under the Unhealthy diet in the male WBM. While the reason behind this dramatic decrease in glucose secretion under the Unhealthy diet has not been reported before, the observed increase for the High Protein diet has been documented. For example, protein feeding in mice has been reported to induce an increase in endogenous glucose production in the kidney and consequently increased glucose release into blood compared to a normal starch diet ^37^. Another striking finding was the significant reduction in pancreatic glucose uptake under the Unhealthy, Average American, and Keto diets compared to other diets in males (**Figure 3C**). The pancreas plays a critical role in blood sugar regulation by secreting insulin. Increased glucose uptake by pancreatic cells is necessary for proper insulin production and secretion. Although, the WBMs do not account for insulin production, these observations suggest a potential link between unhealthy diets high in saturated fat, simple sugars, and processed foods and impaired pancreatic metabolic function.

In females, notable variations in organ contributions to blood glucose levels were observed in the breast and uterus for the Average American and Unhealthy diets, with significant reductions in glucose secretion compared to other diets. The liver also showed decreased glucose secretion under the Keto and Unhealthy diets while the skin and colon experienced diminished glucose uptake from the blood under the Unhealthy, Keto, and Average American diets—further emphasizing the gender-specific responses to diet.

### Pronounced differences in TAG synthesis across diets reflect the influence of high-caloric and fat as well as high-fiber and balanced content

Our analysis of TAG synthesis in adipocytes as surrogate for TAG serum levels revealed pronounced differences across diets (**Figure 4**). In males, the Unhealthy diet exhibited a substantial increase in TAG synthesis compared to other diets, reflecting its high caloric content and unhealthy fats, which contribute to lipid accumulation. This finding aligns with previous studies showing that diets rich in saturated fats and refined sugars promote adipogenesis and lipid storage ^38,39^. Conversely, the Balanced diet exhibited negligible TAG synthesis, highlighting its role in preventing excessive lipid accumulation. The Vegetarian diet also shows slightly reduced TAG synthesis when compared to other diets. This aligns with existing literature suggesting that complex carbohydrate intake —a characteristic of these this diet— promote satiety, reduce calorie intake, and may improve glycemic control by reducing blood sugar spikes after meals, all of which could contribute to decreased TAG synthesis ^40^. In females, the Keto and Unhealthy diets led to the highest while the Balanced, Mediterranean, Vegetarian, Vegan, and DACH diets showed reduced TAG synthesis fluxes. These findings support the adverse impact of high-caloric and high-fat diets and the key role of balanced, fiber-rich diets in regulating TAG storage in adipocytes.

### Differential LDL-C/HDL-C ratios across diets highlight the adverse effects of high-calorie and high-fat diets and the favorable impacts of plant-based diets on lipid profiles

When examining cholesterol metabolism, the analysis of LDL-C/HDL-C ratio provided critical insights. The Unhealthy diet exhibited the highest LDL-C/HDL-C ratio in males (**Figure 5A**), indicating impaired cholesterol metabolism and increased MetS risk. This finding is supported by evidence linking high saturated fat intake with adverse lipid profiles ^41,42,43^). Conversely, the Vegan diet had the lowest (although not significantly lower) LDL-C/HDL-C ratio, reflecting a lower MetS risk. In females, the Keto diet followed by the Unhealthy diet resulted in the highest LDL-C/HDL-C ratios. The high ratio for the Keto diet suggests that this diet adversely affects lipid profiles more severely in females compared to males. Similar to males, the Vegetarian and Vegan diets exhibited the lowest ratios in females (**Figure 5B**). These diets are typically low in saturated fats and high in dietary fibers and unsaturated fats, which promote the synthesis of HDL-C and the clearance of LDL-C from the bloodstream. The low LDL-C/HDL-C ratios for these diets support the potential cardioprotective effects associated with plant-based dietary patterns and their role in reducing the risk factors for MetS ^44^.

### Dietary influence on fatty acid beta-oxidation demonstrates the potential of plant-based diets to rival high-fat regimens in enhancing lipid metabolism

In addition to the four conventional biomarkers implicated in MetS, we also studied the activity of the fatty acid beta-oxidation pathway by focusing on the conversion of butyryl-CoA to acetyl-CoA, the last step in this pathway, as a proxy. While not a conventional marker for MetS, this pathway plays a significant role in lipid metabolism, the dysregulation of which is a known factor in the development of MetS ^27^. In males, the Vegan and Vegetarian diets exhibited the highest fatty acid oxidation activity (**Figure 6A**), likely due to the increased intake of medium-chain fatty acids (MCFAs), which are prevalent in these diets and have been associated with increased fatty acid oxidation ^45^. The Average European Diet followed by the Keto diet also showed high activity. The high fatty acid oxidation activity for the Keto diet was expected due to its high-fat and low carbohydrate content, which shifts energy metabolism towards increased lipolysis and fatty acid utilization. This leads to the production of ketone bodies that provide an alternative energy source for the body when carbohydrate intake is low ^46^. Our findings also suggest that promoting lipolytic activity within the body is not exclusively dependent on the adoption of extreme dietary regimens, such as the Keto diet; rather, a plant-centric diet may offer comparable, or even superior, efficacy. On the other end of the spectrum, the Balanced and Average American diets showed the lowest activity in the male WBM, suggesting a metabolic shift towards other energy sources. In females, we observed divergent patterns where paradoxically the Average American diet, followed by the Balanced, Vegan, and Mediterranean diets had the highest activity, while the Unhealthy and unexpectedly Keto diets exhibited the lowest activity (**Figure 6B**). These results reinforce the effectiveness of plant-based and healthier diets in promoting lipid oxidation. Additionally, the unexpectedly low fatty acid oxidation activity in the female WBM under the Keto diet can be explained by the lower fatty acid oxidation levels in both muscles and heart. This can be attributed to a lower muscle mass and a higher percentage of body fat in women compared to men and inherent gender differences in energy and lipid metabolism ^47^.

### Gender differences in metabolic responses to diets highlight the need for tailored nutritional strategies

Throughout our analysis, we observed several commonalities in the metabolic responses to dietary changes in both males and females as described above, suggesting that these responses are largely independent of gender. However, a number of distinct sex-specific patterns also emerged. For instance, when examining the systemic metabolic response to diet, we noticed that Average American diet clusters with the Average European diet and is proximal to less healthy diets such as the Unbalanced and Keto diets in males (**Figure 2A**); however, it shifts towards the cluster of healthier diets—Vegetarian, DACH, Balanced—in females (**Figure 2B**). Furthermore, the female WBM reveals a rather distinct clustering of healthier diets in relation to MetS biomarkers compared to males (**Figures 2C** and **2D**). Specifically, while some healthy diets like the Mediterranean, Vegan, and DACH aggregate in the lower right quadrant of the t-SNE plot, others (the Balanced and Vegetarian diets) form a separate cluster in the upper right quadrant, indicating a differentiated MetS risk profile—a trend not observed in males.

The overall glucose secretion and TAG synthesis flux are also significantly higher in the female WBM compared to the male. This can be explained by differences in body weight, height, and other physiological parameters used to parameterize the male and female WBMs. These findings also align with recognized sex-specific differences in both glucose metabolism ^48^ and fat storage and utilization ^49, 50^. Additionally, the higher TAG synthesis flux in the female WBM is consistent with slightly higher prevenance of obesity in females compared to males ^51, 52^.

Another notable trend is the greater variation in TAG synthesis levels across different diets in females relative to males, suggesting that men may have a better capacity to regulate fat storage and utilization compared to women. This hypothesis is supported by existing evidence showing significant differences in fat distribution and metabolism between men and women. Specially, men tend to store more fat in the abdominal region, which is more metabolically active and easier to mobilize for energy, while women store more fat in the thighs and buttocks, which is less readily mobilized ^49, 50^. Furthermore, men generally exhibit greater metabolic flexibility, allowing them to switch between burning carbohydrates and fats more efficiently than women, which could contribute to better regulation of fat storage and utilization ^49^.

In regard to lipid metabolism, again, more pronounced variations across diets were observed for LDL-C/HDL-C ratio in the female WBM compared to the male. This aligns with prior reports that women often show greater changes in HDL cholesterol levels, which influence the LDL/HDL ratio, in response to dietary changes such as high fat and high carbohydrate feeding compared to men ^53, 54^. Regarding fatty acid oxidation, the Balanced and Average American diets are associated with the lowest activity in males (**Figure 6A**), while in females, a reversed pattern emerges where these diets exhibit the highest oxidation activity (**Figure 6B**).

Notably, the Keto diet demonstrates divergent trends between genders with regard to the MetS biomarkers. For instance, it induces substantially higher TAG synthesis and elevated LDL-C/HDL-C ratios in females (F**igure 4B**)—a pattern not observed in males. This finding is corroborated by recent research showing that the Keto diet leads to increased adiposity more significantly in female mouse models compared to males as well as sex-specific differences in lipid profiles ^55^. Moreover, while the Keto diet promotes high fatty acid oxidation in males (**Figure 6A**), it shows the second lowest activity in females (**Figure 6B**). This suggests that the Keto diet may not be an effective strategy for weight loss and lowering the risk of MetS in females. This is consistent with prior reports that men tend to experience a more significant increase in fat oxidation after following a Keto diet compared to women ^56^, Sex-specific metabolic responses to the Keto diet has been recognized in recent research ^55–57^, highlighting the significance of our findings.

The Average American diet also presented several counterintuitive results, especially for the female WBM. Notably, unexpectedly low TAG synthesis and storage were observed in both male and female models under this diet (**Figures 4**). Additionally, this diet displayed the highest fatty acid oxidation activity in the female WBM (**Figure 5B**). This unexpected outcome may reflect the diet’s high content of saturated fats and refined carbohydrates, which, despite their association with negative health outcomes, could inadvertently fuel an increase in fatty acid metabolism in females. The low TAG synthesis along with high fatty acid oxidation activity might also indicate a compensatory metabolic mechanism by the body aimed at managing the excessive intake of saturated fats in this diet. Other unexpected metabolic responses to the Average American diet in females, further complicating our understanding of its metabolic impact include one of the lowest LDL/HDL ratios (**Figure 6B**), and abnormal glucose exchange patterns in female organs such as the skin, retina, and adrenal gland (**Figure 2D**). These findings may reflect an unconventional metabolic response and metabolic dysregulation in females that is unique to this diet’s specific nutrient profile. This hypothesis is supported by the unique positioning of the Average American diet in females t-SNE plot (**Figure 2D**). These findings may also hint at limitations of the female WBM.

Collectively, these observations indicate the sex-specific metabolic processing of the micronutrient content in diets and consequently specific metabolic response of males and females to both healthy and unhealthy diets in relation to the MetS risk. Thes also underscores the need for gender-specific dietary recommendations to mitigate MetS risk.

### While WBMs reveal key insights into the impact of diet on MetS risk, their inherent limitations illuminate pathways for future research

It is important to also highlight some important limitations of the present study. Notably, the WBMs lacked the ability to capture metabolite concentrations in the blood as the they are stoichiometric models that can predict reaction fluxes only. Therefore, although the observed trends for the secretion flux of the MetS molecular biomarkers into the blood are expected to correlate with their serum levels, the actual changes in serum concentrations of these metabolites in response to dietary variations were not directly observable. Another critical constraint of our studies is the exclusion of broader physiological regulatory mechanisms such as hormone activity, allosteric regulation of enzymes, signal transduction, and gene regulation, which are not captured within the WBMs. This omission might lead to an incomplete portrayal of the diet’s influence on MetS biomarkers. For instance, most people with MetS also suffer from insulin resistance ^58^ making it more difficult for tissues/organs in the body to respond to insulin and to uptake glucose, but these effects cannot be captured within the WBMs.

These limitations necessitate a cautious interpretation of our findings as they are confined to the specific scope of the male and female WBMs, which focus solely on metabolic pathways and reaction fluxes and exclude non-metabolic regulatory processes and feedback mechanisms that significantly influence physiological responses to dietary inputs. Some predictions by the WBMs might be interpreted in light of these limitations. For example, while both the male and female WBMs predict glucose, LDL-C, HDL-C secretion by adipocytes under the examined diets, current literature does not support direct glucose, LDL-C, and HDL-C secretion by adipocytes into the bloodstream, Adipocytes are, however, recognized to play a pivotal role in maintaining glucose homeostasis and regulating insulin sensitivity ^59,60^. Additionally, they play a key role in the lipid metabolism and cholesterol efflux ^61^, thereby influencing the blood levels of these lipoproteins. The inclusion of glucose, LDL-C, and HDL-C exchange reactions between the adipocytes and blood in WBMs, along with their predicted secretion into the blood could thus reflect an oversimplification of the adipocytes’ role in glucose and lipid metabolism in these models.

Therefore, while our findings are indeed largely corroborated by prior experimental or clinical studies, reinforcing their significance, some of the novel insights uncovered in our analysis warrant further validation in future research. These novel findings offer a strong foundation, guiding further targeted empirical or clinical investigations.

### Conclusion

In this study, we presented a rigorous in silico investigation that contributes to a deeper understanding of the impact of various dietary regimens on the risk of developing MetS. Our findings not only revealed novel insights into the molecular mechanisms underlying the relationships between poor diets and elevated risk of MetS, but also challenge some established notions about diet and metabolic health. Notably, this study underscores the importance of considering the micronutrient composition of diets and gender differences in response to dietary interventions. This aligns with the emerging trend in nutritional science advocating for personalized dietary interventions based on individual metabolic profiles ^62^. Although further empirical research is needed to confirm our novel findings, our study demonstrates the potential of in silico modeling in elucidating intricate biological responses to dietary interventions, which are challenging to parse using in vitro, in vivo, or clinical studies. For instance, by using this approach we were able to computationally identify the contribution of individual organs/tissues toward the metabolic processes necessary for metabolic homeostasis or disruption thereof in MetS. Furthermore, the use of in silico models can mitigate the influence of confounding environmental variables typically seen in dietary intervention studies, thereby offering clearer insights into the metabolic reprogramming associated with MetS. Overall, our findings have the potential to lead to a more comprehensive understanding of metabolic health and diet and to inform more effective dietary intervention strategies to manage or mitigate the risk of MetS.

## Materials and Methods

### Whole-body models

We utilized sex-specific WBMs constructed by Thiele et. al ^11^ and constrained with metabolomic and physiological. The latest versions of the male and female WBMs were obtained from the Virtual Metabolic Human (VMH) database ^19^. The male model captured a total of 81,094 reactions and 56,452 metabolites, representing a typical male subject with a body weight of 70 kg, a height of 170 cm, a consistent heart rate of 67 beats per minute, a stroke volume of 80 mL, a cardiac output of 5360, and hematocrit of 0.400 ^19^. The female model captured a total of 83,521 reactions and 58,851 metabolites, a body weight of 58 kg, height of 160 cm, and with all remaining parameters having the same values as the male model ^19^.

### In silico diets

The in silico diet formulations for the Average European, DACH, High Protein, High Fiber, Mediterranean, Unhealthy, Keto, Gluten Free, Vegetarian, and Vegan diets were obtained from the pre-defined diets section on the VMH database ^19^. The Average American diet is based on the average diet of American males ages 20 and older according to a 2007-2008 NHANES Study and its in silico formulation was obtained from a previous study ^63^. The in silico composition of the Balanced diet was also adopted from Sahoo and Thiele ^64^.

### Computational simulations of WBMs

We performed the computational simulation of the WBMs under different diets by using parsimonious Flux Balance Analysis (pFBA) where the Euclidean norm of reaction fluxes in the model was minimized as the objective function, following that in Thiele et. al ^11^. Other than the bounds on uptake reactions for dietary compounds, the rest of the lower and upper bounds on internal reactions and all the constraints were the same as those in the WBMs ^11^. Computational simulations were conducted in MATLAB 2019b (MathWorks Inc) by utilizing the COBRA (Constraint-Based Reconstruction and Analysis) Toolbox ^65^. We used IBM ILOG CPLEX Optimization Studio V12.10.0, to solve the pFBA optimization problem. All simulations were performed on a local computer with a 3.3 GHz Dual-Core Intel i7 processor and 16 GB of memory.

### Dimension reduction techniques

We conducted dimension reduction analyses using MATLAB 2019b’s PCA and t-SNE functions (with default parameters for t-SNE), which are part of the Statistics and Machine Learning toolbox for MATLAB.

## Declarations

### Availability of data and material

All data from this study is available in the supplementary files.

### Authors’ contributions

ARZ conceived the study and, together with DSA, interpreted the results and drafted the manuscript. DSA performed all analyses. CVM assisted with computational simulations and provided the in silico Average American diet formulation. All authors have read and approved the final manuscript.

## Acknowledgements

The authors gratefully acknowledge Dr. Carol Ehrlich and Dr. Marouen Ben Guebila for insightful feedback on this work.

## Declaration of generative AI and AI-assisted technologies in the writing process

During the preparation of this work the author(s) used the Microsoft Copilot for literature search and OpenAI’s ChatGPT in order to refine the language in certain sections of the manuscript. After using this tool/service, the author(s) reviewed and edited the content as needed and take(s) full responsibility for the content of the publication.

## Supplementary files

**Supplementary File 1**. The file contains the micronutrient compositions of the diets, list of reactions related to the MetS biomarkers and fatty acid beta-oxidation pathway within the male and female WBMs, along with their predicted flux values under the examined diets.

